## Supplementary material for "aPKC and F-actin Dynamics Promote Hippo Pathway Polarity in Asymmetrically Dividing Neuroblasts": Suppl. Figs. 1-2 and list of suppl. moviess

**Figure S1**

**A**

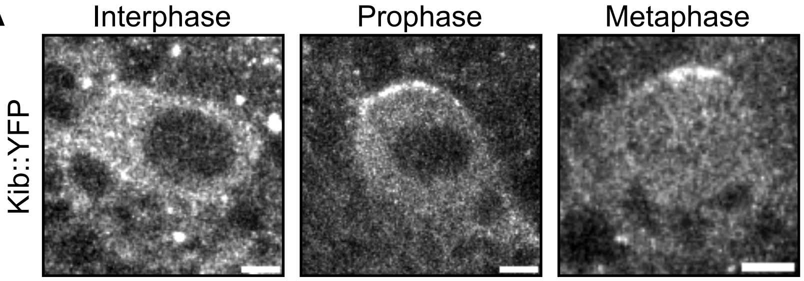


**Endogenously-expressed Kibra polarizes in living neuroblasts. A.** Interphase, prophase, and metaphase images of neuroblasts endogenously expressing Kib::YFP.

Figure S2

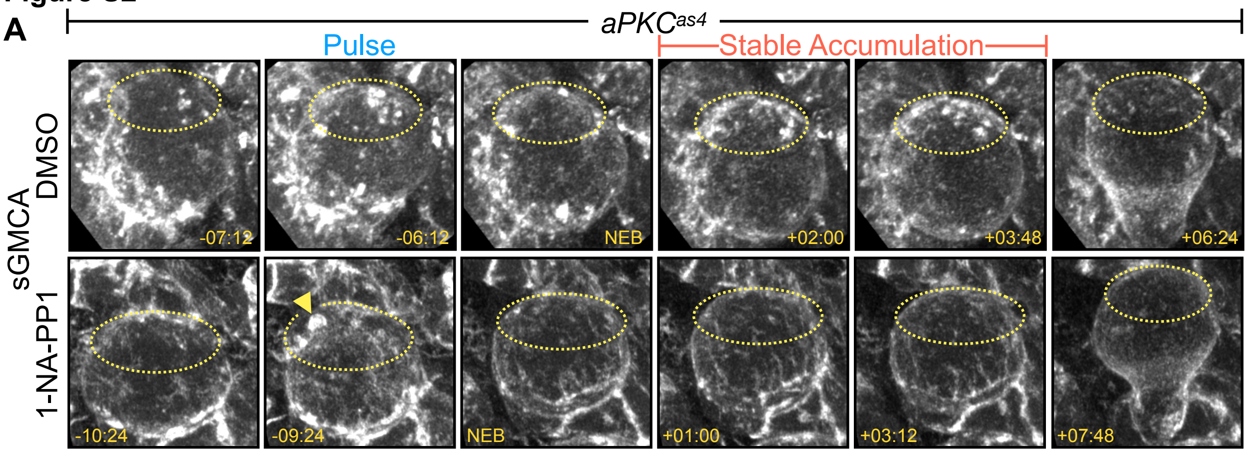


**aPKC regulates Actin dynamics in live neuroblasts. A.** Mitotic neuroblasts expressing the F-actin reporter sGMCA (*spaghetti squash* driven, moesin-alpha-helical-coiled and Actin binding site fused to GFP) from *aPKC*$^{as4}$ animals treated with DMSO as a control or the *aPKC*$^{as4}$ inhibitor 1-NA-PP1. Yellow dotted lines approximately demarcate the apical cortex. Yellow arrowhead indicates an example of a cortical bleb. Pulse refers to a single frame in which short-lived F-actin accumulation at the apical cortex was captured. Stable Accumulation refers to multiple frames in which long lived F-actin accumulation at the apical cortex was captured. Note the cortical Actin blebbing during the Pulse phase and diminished apical accumulation during the Stable Accumulation phase in the treated neuroblasts.

**Movie S1 (related to Fig. 1B).** Mitotic neuroblast expressing Ubi>Kib-GFP.

**Movie S2 (related to Fig. 1C).** Super-resolution view of the apical cortex of a mitotic neuroblast expressing Ubi>Kib-GFP in prophase.

**Movie S3 (related to Fig. 2A).** Mitotic neuroblast coexpressing aPKC-Halo and Ubi>Kib-GFP.

**Movie S4 (related to Fig. 2B).** Mitotic neuroblast coexpressing Baz-GFP And Ubi>Kib-Halo.

**Movie S5 (related to Fig. 2C).** Mitotic neuroblast coexpressing Dlg-GFP and Ubi>Kib-Halo.

**Movie S6 (related to Fig. 3A).** Effect of acute aPKC inhibition with 1-NA-PP1 on Ubi>Kib-GFP polarization in *aPKC*$^{as4}$ neuroblasts.

**Movie S7 (related to Fig. 4B).** Mitotic neuroblast expressing Ubi>Kib$^{DaPKC}$-GFP.

**Movie S8 (related to Fig. 4C).** Mitotic neuroblast expressing Ubi>Kib$^{858\text{-}1288}$-GFP.

**Movie S9 (related to Fig. 5A).** Effect of F-actin inhibition with Latrunculin A on aPKC-Halo polarization. Same neuroblast as Supplemental Movie 10.

**Movie S10 (related to Fig. 5B).** Effect of F-actin inhibition with Latrunculin A on Ubi>Kib-GFP polarization. Same neuroblast as Supplemental Movie 9.

**Movie S11 (related to Fig. 6A-C).** Apical max projection of a mitotic neuroblast coexpressing LifeAct-Halo and Ubi>Kib-GFP. Top-down view.

**Movie S12.** Mitotic neuroblast expressing Ubi>Mer-Halo. This view is of a single cortical slice.

**Movie S13 (related to Fig. 7A).** Mitotic neuroblast expressing Ubi>Sav-GFP.

**Movie S14 (related to Fig. 7A).** Effect of Kib knockdown with insc>Kib-RNAi on Ubi>Sav-GFP polarization.

**Movie S15 (related to Fig. 7B).** Effect of Sav knockdown with insc>Sav-RNAi on Ubi>Kib-GFP polarization.

**Movie S16 (related to Fig. 8A).** Mitotic neuroblast expressing GFP::Wts.

**Movie S17 (related to Fig. S2A).** Effect of acute aPKC inhibition with 1-NA-PP1 on cortical F-actin dynamics, visualized with the F-actin reporter sGMCA (*spaghetti squash* driven, moesin-alpha-helical-coiled and Actin binding site fused to GFP).
